# An Agent-Based Model of Protein Polymerization Dynamics: Focus on the Actin System

**DOI:** 10.64898/2026.01.02.697380

**Authors:** Riccardo Tarantino, Salvatore Contino, Livia Gugliotta, Giuseppe Indelicato, Greta Panunzi, Giorgio Bertolazzi, Valentino Romano

**Author notes:** These authors contributed equally to this work (Riccardo Tarantino, Salvatore Contino).

## Abstract

Actin polymerization is a critical cellular process involved in a wide range of activities, from cell motility to cytokinesis. The complex behavior of this molecular system, resulting in three different phases (i.e., nucleation, elongation, and steady state) is clear by looking at the way these dynamics emerge from a large number of interactions between different proteins, regulatory elements, and signaling pathways. In this article, we present an agent-based model of actin polymerization dynamics implemented with the NetLogo simulation platform and focus on the time evolution of actin filaments length distribution in two dimensions starting from a pool of free G-actin monomers. Stochastic simulations were able to reproduce the three main steps of the in vitro process, i.e., nucleation, extension, and steady state. In the steady state phase, simultaneous treadmilling of subpopulations of F-actin filaments spontaneously occurs as an emergent property of the system, as shown by several diagnostic criteria. We also found that during the early phases of polymerization the dynamics are characterized by the competition between the formation of new polymers and the elongation of pre-existing ones. In this regard, we highlighted that the balance between nucleation and elongation is particularly sensitive to variation in the initial number of free monomers available for association, as well as on other parameters included in the model. Altogether, our observations are consistent with the experimental findings and confirm the satisfactory performance of the model, as well as its ability to simulate complex mechanisms by manipulating a few parameters.

## Introduction

Actin (42 kDa) is a highly conserved protein expressed in several cell types in a wide range of organisms, from bacteria to humans [1,2]. Actin polymerization is a fundamental process that occurs in cells, where it plays a crucial role in many activities, including cell motility, cell division, intracellular transport, and maintenance of cell shape. This process involves the assembly of actin monomers (G-actin) into long, filamentous structures called actin filaments (F-actin). Studies performed in cell-free systems have been instrumental in advancing our understanding of the molecular mechanisms and dynamics of actin polymerization [3]. Thanks to these studies, we know that the process can be divided into three main stages:

1. Nucleation: actin polymerization begins with the nucleation phase, where a small pool of actin monomers forms the initial nucleus for filament growth. Nucleation can occur spontaneously [4,5];
2. Elongation: once the critical nucleus is formed, actin monomers can be added to both ends of the filament in a process called elongation. Actin monomers have a polar structure, with a fast-growing or “barbed” end (also known as the plus end) and a slow-growing or “pointed” end (also known as the minus end). The addition of actin monomers is more favorable at the barbed-end, resulting in rapid elongation at that end, while the pointed-end remains relatively stable[1,2];
3. Steady state: in the steady state phase, actin filaments continue to grow at the barbed-end by adding monomers, while simultaneously undergoing dissociation or depolymerization at the pointed-end. This dynamic behavior is known as treadmilling, where actin monomers are constantly exchanged between the filaments and the cytoplasm, maintaining a relatively constant filament length [6].

Various proteins in cells assist and regulate the process of actin polymerization. These proteins play critical roles in nucleating actin filaments, promoting filament elongation, capping filament ends, and facilitating filament severing or disassembly. These are just a few examples of the many proteins involved in actin polymerization. The coordinated actions of these proteins, along with additional actin-binding proteins, regulatory factors, and signaling pathways, tightly regulate actin dynamics and allow for the precise control of actin filament assembly, disassembly, and organization within cells [4,5,7,8].

Mathematical and computational modeling have made significant contributions to our understanding of the actin polymerization process by providing quantitative frameworks to explain and predict several aspects of actin dynamics. For example, several models have been developed to describe the rates and kinetics of actin polymerization. These models incorporate the steps involved in the polymerization process and provide equations that describe the changes in filaments length and the concentration of actin monomers over time [9–12]. By fitting the outcomes of such models to experimental data, researchers can estimate parameters such as the critical concentration for filaments growth and the rate constants for polymerization and depolymerization. Moreover, some other models have helped in explaining the phenomenon of actin treadmilling. Indeed, by incorporating the rates of monomeric addition and dissociation at both ends of the filaments, they have provided insights into the factors that regulate treadmilling and filament lengths at the steady state. In particular, they have improved our knowledge of how changes in the concentrations of actin-binding proteins and nucleotides affect the dynamics of treadmilling [13].

A particular class of computational models is represented by agent-based models (ABMs). The fundamental principle of agent-based modeling is that the aggregate behavior of any natural, social, or engineering system of interest can be reconstructed from the bottom up by simulating the individual behaviors performed by the subunits that “populate” a lower level of the system itself. In agent-based modeling jargon, these subunits are called “agents”. More specifically, agents are autonomous computational individuals that have certain properties or state variables and execute some activities indicated by the programmer through algorithms. If the conditions imposed by the programmer are met during a computer run, agents enact the behavior described in these algorithmic instructions. ABMs have been shown to be especially powerful in exploring the dynamics of complex systems [14]. Complex systems are usually characterized by feedback loops, non-linear events, stochasticity, and the presence of global patterns (or *emerging properties*) which spontaneously arise from the dense network of interactions between the lower-level components of the system. Considering the bottom-up determination of cellular outcomes and states, which can be regarded as emerging properties generated by a very large number of interacting molecular entities, actin polymerization too can be viewed and studied as a complex system, especially in light of the many feedback loops among these entities.

Although there are already several studies that have investigated a variety of aspects of actin dynamics using ABMs [15–18], to the authors’ knowledge, the present model is the first implemented in the NetLogo simulation platform [19] that is able to reproduce actin polymerization dynamics. NetLogo is both a programming language with a simple syntax and a modeling environment which was specifically designed for the implementation of agent-based models, also including a customizable interface through which users can interact with simulations and observe the dynamic behavior of agents (e.g., viruses, cells, vehicles, investors, and so on) whose activities are regulated by an algorithm. For a more detailed examination of agent-based modeling and NetLogo, see [14] and [20]. NetLogo enabled us to reproduce the three main steps of polymerization, as well as global treadmilling and the competitive interplay between nucleation and elongation of polymers [21,22], in terms of interactions between agents, i.e., actin monomers whose role changes over time or in response to variation in some key parameters.

The bottom-up generation of these emerging patterns, together with the ability to fully preserve their inherent stochastic component, are among the main strengths of the agent-based modeling approach we adopted. Furthermore, NetLogo enables wide usability also by researchers who are unfamiliar with computational models, due to its user-friendly Interface and captivating visual component, plus the possibility of automatically running large sets of simulations by using its integrated tool BehaviorSpace. This option is highly valuable, as it provides a wealth of data for statistical analysis in a relatively short time and in a context where the main variables are easily controlled by the scientist.

In summary, our main aim is to enhance the understanding of all phases of actin polymerization, highlighting the relationship between the individual behavior of the molecular actors involved and the global properties of the system, especially treadmilling and the balance between nucleation and elongation. Indeed, while all these dynamics are easy to observe in the NetLogo model, they are instead more elusive both in vitro and in vivo. In the following sections, we present the design and parameters of our model (Materials and methods), report several statistical analyses aimed at testing the general performance of the system and confirming its validity from a qualitative point of view (Results), discuss our observations (Discussion), and comment on some possible future extensions (Conclusions).

## Materials and methods

### Model overview and conceptual framework

“Actin Polymerization” is a stochastic model aimed at reproducing the three stages of actin polymerization in vitro starting from an initial pool of G-actin monomers: nucleation, extension, and steady state. More specifically, in the steady state phase, sub-populations of polymers undergo treadmilling (see Results section). The version of NetLogo used in this study is version 6.2.0, freely downloadable from the following url: https://ccl.northwestern.edu/netlogo/6.2.0/. The model’s interface, which contains the simulation world and several items used to adjust parameter values and display the outputs of runs, is shown in Fig. 1.

**Fig. 1.**
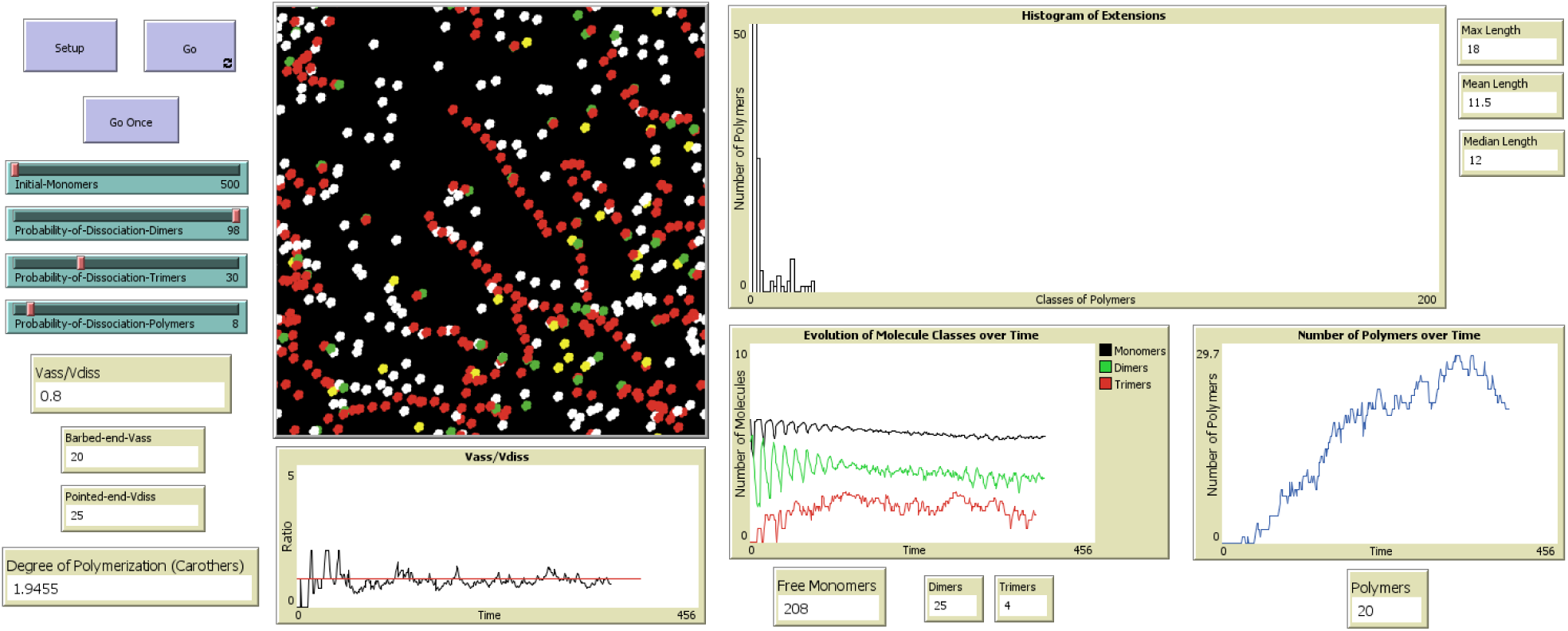
Interface of the NetLogo model “Actin Polymerization”. In this simulation, we used a very small number of agents to improve the visual component of the digital system.

The agents which appear at the beginning of each simulation are free monomers (white color), whose number is regulated by the parameter *Initial-Monomers*. Free monomers move randomly in a two-dimensional, toroidal space. When they are available for polymerization (see variables *Receptivity* and *Restoration* in Table 1), they behave as G-actin-ATP monomers [1]. When two free, available G-actin-ATP monomers share the same coordinates on NetLogo lattice, they produce an unstable dimer and, soon after, decide whether to form a stable dimer or dissociate into the two original monomers (i.e., the nucleation phase starts). The probability of formation of a new, stable dimer is regulated by the parameter *Probability-of-Dissociation-Dimers*. The same mechanism is applied to trimers: when a free, available G-actin-ATP monomer and the barbed-end (red color) of a stable dimer share the same coordinates, they form an unstable trimer, which can eventually become a stable trimer or release a free monomer and a dimer, based on the parameter *Probability-of-Dissociation-Trimers*. Dimers and trimers are very unstable and tend to dissociate rapidly [23], although the model allows to regulate this degree of instability. We can assume that the regulation of *Probability-of-Dissociation-Dimers* and *Probability-of-Dissociation-Trimers* may correspond either to the presence/absence of nucleators (e.g., Formin and Arp 2/3), or to their variable activity [24]. In particular, very high values of these probabilities may indicate the absence or almost absence of such nucleators. Vice versa, low values of these probabilities would correspond to the presence of nucleators. Monomers released by dimers and trimers are reused in new nucleation cycles, after a short period of non-receptivity for further associations. This time interval corresponds to the value of the counter *Restoration*, whose function may be associated with the provisional effects of binding of the protein Profilin to monomers [25].

**Table 1.** Parameters, state variables, and interface items included in the NetLogo model.

| **Parameter/state variable/item** | **Description** |
| --- | --- |
| *Time* | Simulation time expressed in ticks, i.e., NetLogo time steps |
| *Initial-Monomers* | Total number of free, available monomers at the beginning of the simulation |
| *Free Monomers* | Total number of free monomers in the current tick |
| *Dimers* | Total number of dimers in the current tick |
| *Trimers* | Total number of trimers in the current tick |
| *Polymers (P)* | Total number of polymers in the current tick (i.e., all filaments from tetramers above) |
| *Extension* | Number of monomeric subunits of a polymer |
| *Polymer-Number* | ID number assigned to each polymer |
| *Max length* | Extension of the longest polymer in the current tick |
| *Mean length ($L_m$)* | Mean extension of polymers in the current tick |
| *Median length* | Median extension of polymers in the current tick |
| *Receptivity* | Availability of free monomers for further associations. It is a binary value, i.e., either “receptive” or “non-receptive” |
| *Restoration* | Time in ticks required to restore the *Receptivity* of a monomer (fixed at 10 ticks) |
| *Probability-of-Dissociation-Dimers* | Regulates the probability that each unstable dimer dissociates into two free, non-receptive monomers |
| *Probability-of-Dissociation-Trimers* | Regulates the probability that each unstable trimer dissociates into a free, non-receptive monomer and an unstable dimer |
| *Probability-of-Dissociation-Polymers* | Regulates the probability that a pointed-end dissociates from a polymer. It is applied at each tick |
| *Barbed-end-$V_{ass}$* | Monomers association rate at the barbed-end, i.e., the number of barbed-ends of polymers in the current tick |
| *Pointed-end-$V_{diss}$* | Monomers dissociation rate at the pointed-end, i.e., the number of dissociations of pointed-ends from polymers in the current tick. This value is maintained until the dissociated monomer associates again with an oligomer or a polymer |
| *$V_{ass} / V_{diss}$* | Association rate/Dissociation rate ratio |
| *Degree of Polymerization (Carothers)* | Result of the Carothers Equation (see text) in the current tick |

Starting with the trimer, again by addition to the barbed-end, polymer synthesis begins (elongation phase). The shortest stable polymer in the model is the tetramer [26,27]. We easily identify each polymer and its length in terms of monomeric subunits (*Extension*) through its ID number, that is, *Polymer-Number* (see Table 1). As for oligomers and polymers movement, all intermediate subunits and barbed-ends follow the random movement of the pointed-end, which acts as a “leader” in a snake-like configuration. Although it does not accurately represent the shape and motility of real actin filaments of limited length, this pattern of movement does not affect polymerization dynamics in any way, even in cases where subunits of a filament appear to overlap with other subunits. Fig. 2 shows what a single filament of short length (i.e., 30 monomeric subunits) looks like during a simulation.

**Fig. 2.**
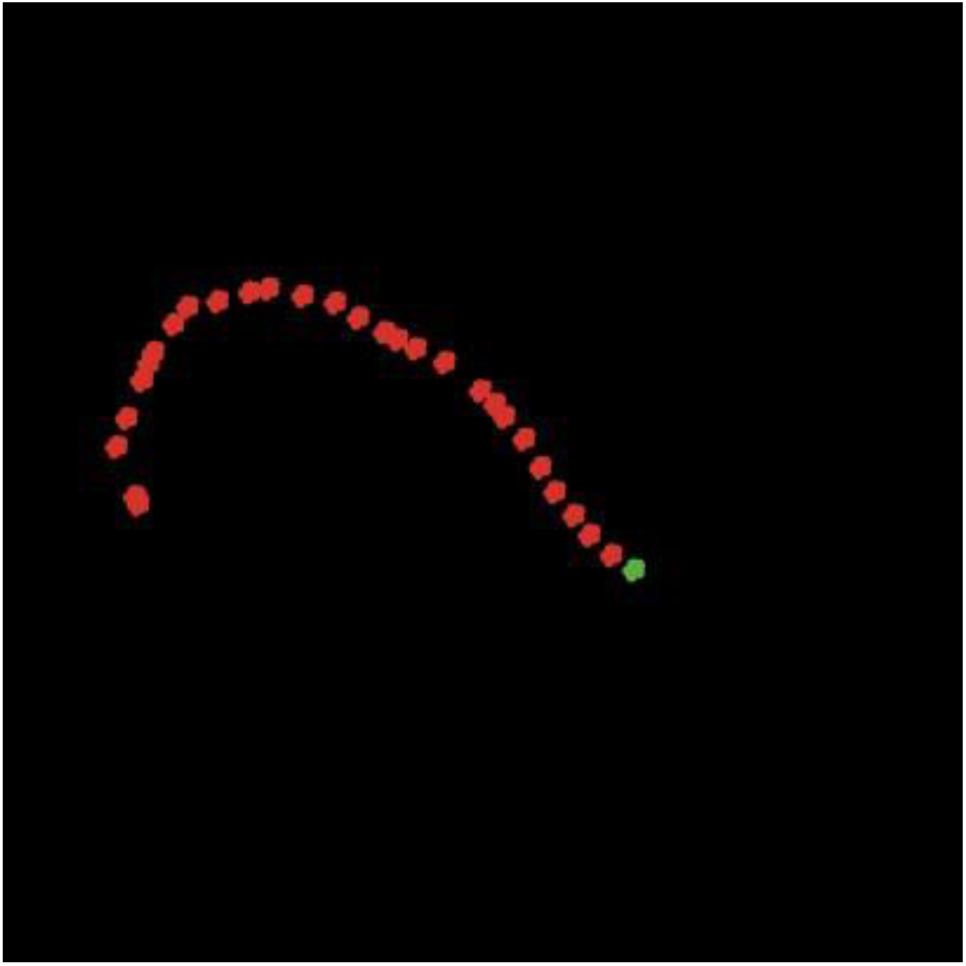
An actin filament in the model. Shape of a single actin filament during a simulation.

Dimers, trimers, and all higher-order oligomers and polymers are oriented and have a well-defined polarity, meaning that they always have a barbed-end (red color, that is also the color of all intermediate monomeric subunits) and a pointed-end (green color). Furthermore, their polarity reflects the direction in which polymerization occurs: while the association of new subunits always occurs from the barbed-end, dissociation occurs from the pointed-end [1,2]. Consequently, each polymer loses a monomer from the pointed-end with a probability which is calculated at each time step (i.e., a NetLogo tick) and corresponds to the selected value of the parameter *Probability-of-Dissociation-Polymers*, whereas each new monomeric subunit is added to the barbed-end by co-localization. The unidirectionality of this phenomenon in the model is a simplification of the real biological pattern. Dissociation of monomeric subunits from the pointed-end determines a reorganization of roles in that polymer (i.e., the intermediate subunit following the previous pointed-end becomes the new pointed-end of the filament). Each monomer that has just dissociated from a filament is yellow and temporarily unavailable for further associations (*Receptivity* = “non-receptive”). For low values of the parameter *Probability-of-Dissociation-Polymers*, we simulate the spontaneous process of dissociation due to the conversion of G-ATP to G-ADP [1]. For higher values of this parameter, we simulate the activity of Cofilin, an actin regulatory protein promoting depolymerization [28]. When an equilibrium between the association and dissociation of monomers in filaments is reached, the system is in the steady state phase.

All parameters, main state variables, and interface items implemented into the model are reported in Table 1. For a visual representation of the processes executed by the model at each time step, see Fig. 3.

**Fig. 3.**
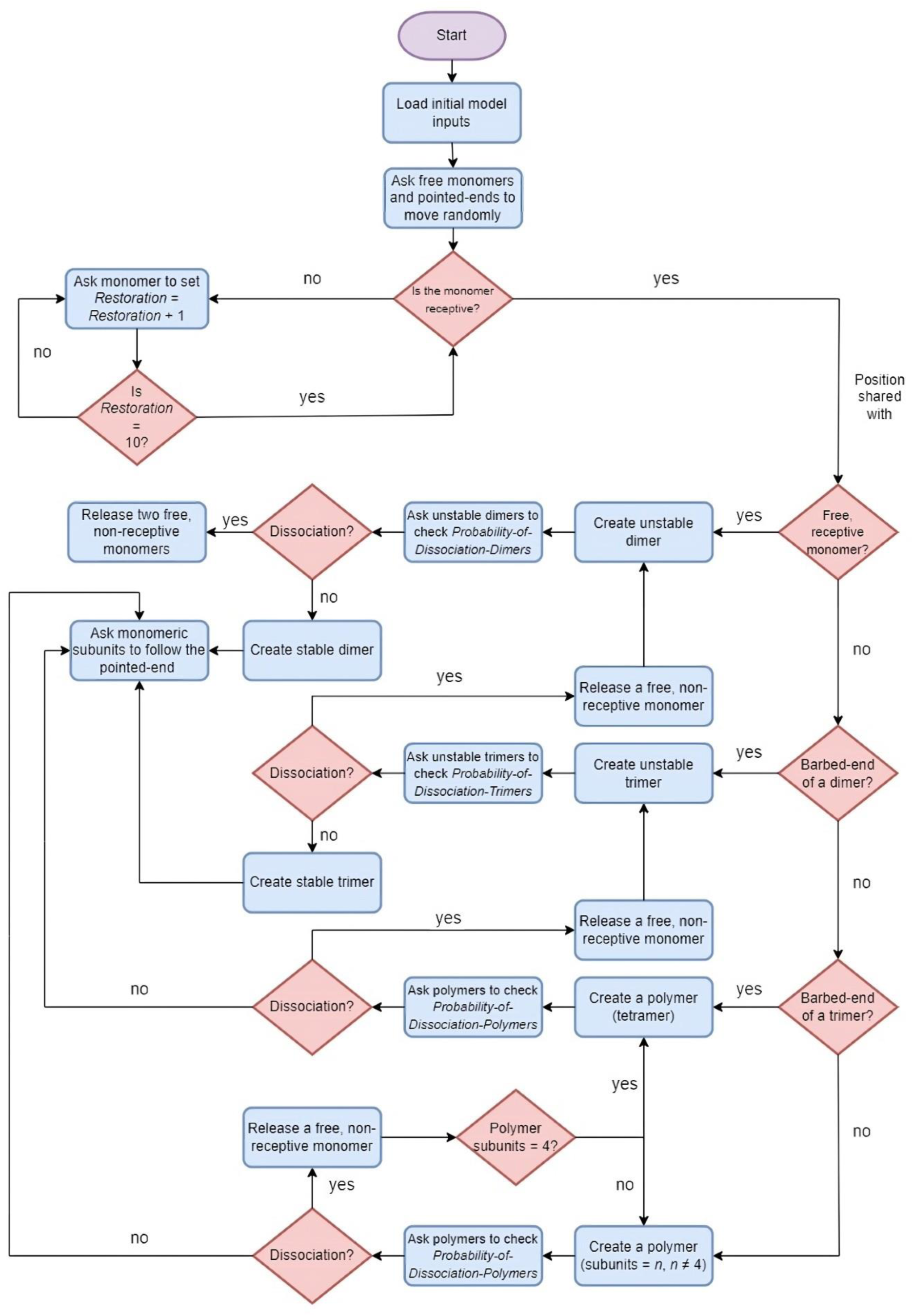
Flowchart of the algorithm. This set of instructions is reiterated at each NetLogo tick (i.e., a discrete time step).

We also compiled a list of similarities and differences between our model and the corresponding phenomena observed in vitro (Table 2).

**Table 2.** Comparison between the actin polymerization in vitro system and the “Actin Polymerization” NetLogo model.

| **N** | **In vitro system** | **“Actin Polymerization” NetLogo model\*** |
| --- | --- | --- |
| 1 | 3D space in a test tube | 2D space (NetLogo lattice) |
| 2 | Number of monomers > 1,000,000 | Maximum number of monomers = 50,000<br>(see comment 1 below) |
| 3 | **Dimers, trimers, and polymers are oriented (i.e., barbed-end and pointed-end) [1,2]** | **Dimers, trimers, and polymers are oriented** |
| 4 | **Free monomers, dimers, trimers, and polymers move with random walk** | **Free monomers, dimers, trimers, and polymers move with random walk** |
| 5 | Time in seconds | Time in NetLogo ticks |
| 6 | Presence of salts, $\text{Ca}^{++}$ and $\text{Mg}^{++}$ in the solution | Salts, $\text{Ca}^{++}$ and $\text{Mg}^{++}$ were not implemented |
| 7 | **Presence of G-actin-ATP monomers and of G-actin-ADP-monomers [1]** | **Presence of G-actin-ATP monomers and G-actin-ADP-monomers was implemented by the *Receptivity* variable (see comment 2 below)** |
| 8 | $K_{\text{ass}}$ and $K_{\text{diss}}$ estimated for dimers and trimers | Not implemented |
| 9 | **Formation of dimers and trimers [23,24]** | **Dimers and trimers are formed by co-localization in the NetLogo lattice** |
| 10 | **High instability of dimers and trimers [23,24]** | **The degree of instability of dimers and trimers depends on probabilistic procedures and is selected by the user (see comment 3 below)** |
| 11 | $K_{\text{ass}}$ and $K_{\text{diss}}$ estimated for polymers | Not implemented (see comment 4 below) |
| 12 | **The shortest polymers contain four monomeric subunits (tetramers) [26,27]** | **The shortest polymers contain four monomeric subunits (tetramers)** |
| 13 | F-actin polymers are arranged in a double helix | Not implemented. The difference between the model and reality is limited to the visual component |
| 14 | **Polarity of F-actin filaments [1,2]** | **Association occurs from the barbed-end (red) and dissociation from the pointed-end (green)** |
| 15 | On average, F-actin filaments contain up to several thousands of monomers | F-actin filaments have lengths $\leq 200$ monomeric subunits (see comment 1 below) |
| 16 | Presence of further mechanisms such as capping, severing, and annealing | Absence of mechanisms such as capping, severing, and annealing |

\* **Bold text** = similarities, normal text = differences.

1. The dissimilarities listed in items 2 and 15 largely depend on limits to the maximum number of agents that can be used in a simulation, due to the CPU and RAM of the computer used. As a matter of fact, a reduced number of initial monomers also has negative effects on filaments length (see item 15);
2. As for item 7, while it is true that specific subsets of agents corresponding to G-actin monomers of different species were not implemented in the model, the *Receptivity* property in the code allows G-actin-ATP to be distinguished from G-actin-ADP. Indeed, the related *Restoration* variable defines the time interval (in NetLogo ticks) required for a monomer that has just detached from the filament (G-actin-ADP, non-receptive) to be reused for another polymerization cycle, thus becoming G-actin-ATP (receptive). *Receptivity* is also used to indicate the period of unavailability of monomers coming from dissociations of unstable dimers and trimers for further nucleation events;
3. Item 10 describes the high instability of dimers and trimers, which, from in vitro studies, appears to be several orders of magnitude greater for dimers than for trimers. In the simulations made with NetLogo, this difference was approximated by making the two dissociation probabilities equal, although both very high (98%), in order to represent the high instability of both oligomers, whereas greater stability is only found in species with a higher number of subunits, starting with tetramers [8] (see more on the rationale behind the choice of 98% dissociation probability for both dimers and trimers in the Discussion);
4. Item 11 refers to the equilibrium constants for G-actin-ADP and G-actin-ATP related to the binding or detachment of these monomers at the barbed and pointed-ends, respectively. In our model, the different affinities of these monomers for the two ends (reconstructed from in vitro studies) were implemented as follows: the binding of G-actin-ATP moving by random walk occurs only if it co-localizes with the barbed-end of the filament. At the pointed-end, the detachment of a monomer (G-actin-ADP) is controlled by a probability-dependent stochastic procedure, which is regulated by the user. Furthermore, the model keeps track of the rates of association and dissociation at the barbed-end and pointed-end and their ratio. In particular: (1) if V_ass_ > V_diss_ then polymerization prevails, (2) if V_ass_< V_diss_ then depolymerization prevails, (3) if V_ass_ = V_diss_ then treadmilling is taking place. Thus, like the corresponding in vitro system, the three conditions allow identification of whether the number of free monomers is above (V_ass_ > V_diss_), below (V_ass_ < V_diss_) or equal (V_ass_ = V_diss_) to the critical number (not concentration) of monomers. The critical number referred to here is the total number for the whole system, not the number for either end of the filament.

### Simulations

Preliminary analyses were conducted aimed at exploring a wide range of initial conditions and settings, including some extreme conditions (e.g., inhibited dissociation of monomeric subunits from the pointed-ends), with an average of ten simulations for each parametrical investigation. In most of the tests presented here, we used parameter values inspired by biochemical knowledge of the actin polymerization process, although these values were not calibrated using quantitative data (i.e., we tested the model qualitatively and used it as a more general toy model of protein polymerization). For instance, we considered the high degree of instability of both dimers and trimers and the presence of a large number of G-actin-ATP monomers. More specifically, except where indicated, the outcomes reported in the Results were obtained using the following initial conditions: 50,000 *Initial-Monomers*, *Probability-of-Dissociation-Dimers* and *Probability-of-Dissociation-Trimers* = 98%, and *Probability-of-Dissociation-Polymers* = 8%. All parameter values used in our experimental scenarios are also summarized in Table 3.

**Table 3.** Parameter values used in each experimental scenario.

|  | *Initial-Monomers* | *Probability-of-Dissociation-Dimers* | *Probability-of-Dissociation-Trimers* | *Probability-of-Dissociation-Polymers* |
| --- | --- | --- | --- | --- |
| The three steps of actin polymerization | 50,000 | 98% | 98% | 8% |
| Effects of nucleators on the start of elongation | 5,000 | 98% - 50% | 98% - 50% | 8% |
| Evolution of filament length distribution | 50,000 | 98% | 98% | 8% |
| Criteria of detection of treadmilling | 5,000 | 98% | 98% | 8% |
| Effects of dimers dissociation on the steady state | 15,000 | 98% - 30% (during the simulation) | 98% | 8% |
| Competition analysis | From 1,000 up to 40,000 | 98% | 98% | 8% |

### Data mining and statistical analyses

Data mining and statistical analyses (including Kolmogorov-Smirnov test, variance, and density plots) were applied to data exported from NetLogo. In particular, Kolmogorov-Smirnov’s test and variance calculation were applied to follow changes in the type and number of length classes over time (measured in NetLogo ticks). The analysis was performed using the R 4.3.0 software [29].

### Degree of polymerization (Carothers’ equation)

To monitor the average degree or intensity of actin polymerization, we used Carothers’ equation. Organic chemists often define by this equation either the average degree of polymerization of a given polymer or the rate of conversion of monomers to polymers in polymeric synthesis reactions. In our case, this parameter was used to determine whether an increase in the degree of polymerization values (C) reflects an increase in the number of polymers or an increase in polymers length. The following equation was used to calculate the degree of polymerization [30–32]:

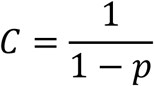

where

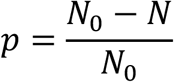

where N_0_ is the number of initial monomers and N is the sum of monomers, dimers, trimers, and polymers.

## Results

### Simulations reproduce the three phases of actin polymerization

The first, relevant outcome observed in all simulations was the emergence of the three phases of actin polymerization: (1) an equilibration phase of monomers, dimers and trimers that would coincide with the nucleation phase described in vitro, (2) an extension phase in which polymers are formed, and (3) a steady state phase in which the number of polymers is constant and there is a slow redistribution of polymer lengths over time. Each of these phases occurs within a given time frame (in ticks) but, while the initial conditions remain the same, the duration of each phase changes from simulation to simulation due to stochastic effects. The three stages of polymerization reproduced in NetLogo are shown in Fig. 4.

**Fig. 4.**
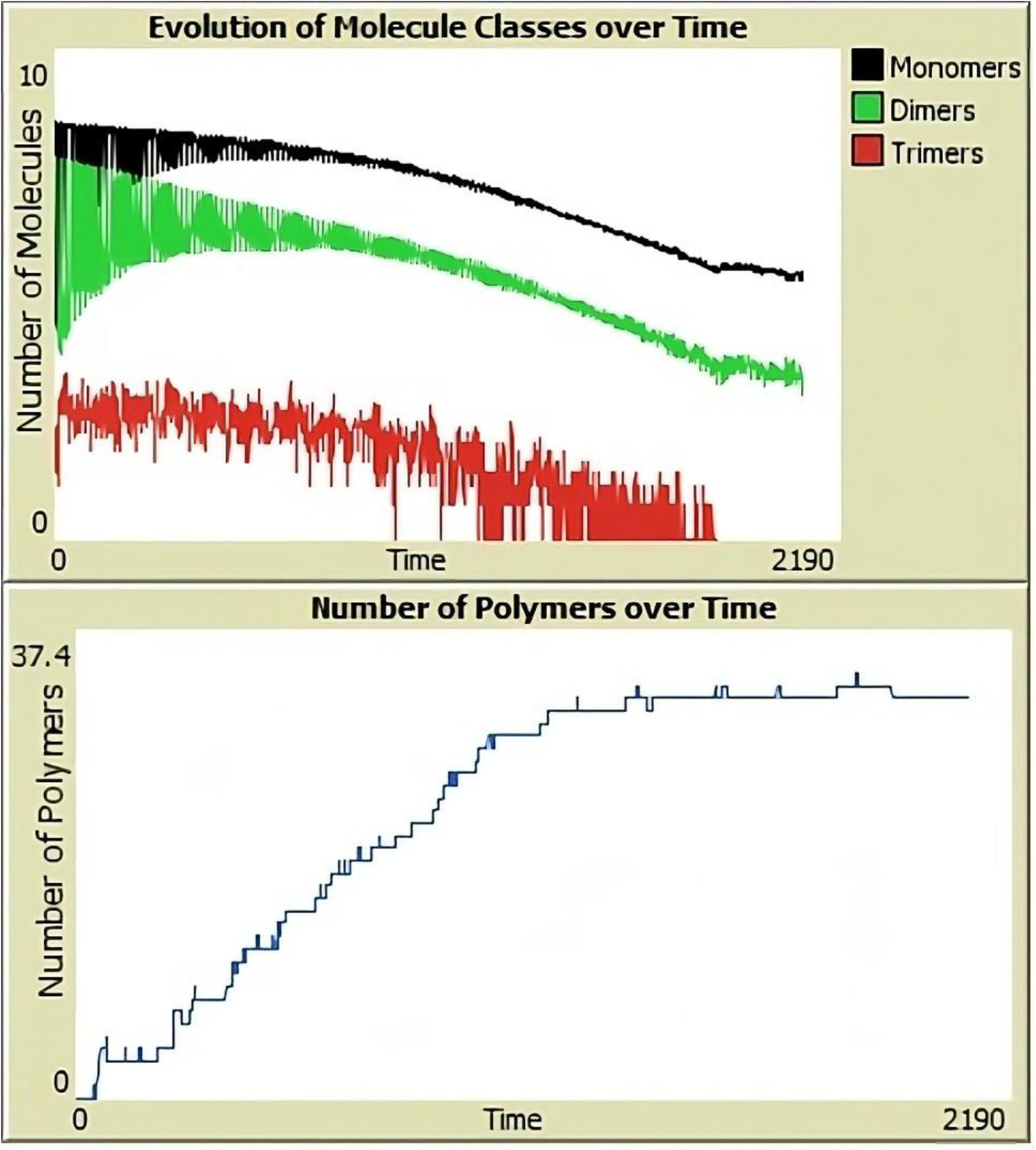
Steps of actin polymerization in the NetLogo model. The plots show the curves corresponding to the change in the number of monomers, dimers, trimers, and polymers during the three stages of polymerization. In the ordinate of the first plot, a logarithmic scale was used for the number of monomers, dimers, and trimers, whereas in the second plot a non-logarithmic scale was used for the number of polymers (the box in the upper right-hand corner displays the color code used for each curve).

A detailed description of each phase is indicated hereafter:

1. During the nucleation phase, there is a greater fluctuation in both the number of monomers (black curve) and the number of dimers (green curve), and less fluctuation in the number of trimers (red curve). Regardless of differences between dimers and trimers’ curves, all fluctuations reflect the thermodynamic instability of these oligomers during this phase [33]. The number of ticks relative to the duration of this phase varies in relation to the number of initial monomers;
2. During the extension phase the curve of polymers (in blue) increases whereas the curves of monomers, dimers, and trimers (in black, green, and red, respectively), decrease gradually as polymers’ curve increases. While the formation of new filaments from dimers and trimers reduces the quantity of these oligomers, the gradual consumption of monomers is caused by the prevalence of their association with filaments over dissociation [33]. The number of ticks relative to the duration of this phase varies according to both the number of initial monomers and the probability value of polymers dissociation;
3. During the steady state phase, we observed that: (1) the polymers’ curve (in blue) reached a plateau and stabilized itself, (2) the curves of the number of monomers (in black) and dimers (in green) decreased until they stabilized themselves at low values (never reaching zero), and (3) trimers’ curve (in red), on the other hand, decreased until it disappeared permanently (few trimers or none). This stabilizing behavior has also been observed in vitro, where simultaneous growth of filaments at one end and shortening at the other end keep the number and length of polymers unchanged while maintaining the concentration of monomers constant [13]. The number of ticks required to reach the steady state varies according to both the number of initial monomers and the probability of polymers dissociation. This dynamic equilibrium phase is the most prominent step of actin polymerization, since it enables the cell to preserve its structural and functional organization. Indeed, the steady state confers cytoskeletal networks a high degree of plasticity, that is necessary to allow cells to adapt to external changes [34]. However, it is important to stress that the mentioned steady state is to be intended as a statistically stationary phase, that is a steady phase characterized by intermittent, transient phenomena (see the Discussion section).

We were also interested in simulating the systemic effects of different classes of nucleators acting on dimers and trimers (reviewed in [35]). We collected data on the time (in ticks) required for the transition between nucleation and elongation. Runs involving a fixed number of *Initial-Monomers* (5,000), fixed *Probability-of-Dissociation-Polymers* (8%), and variable values of *Probability-of-Dissociation-Dimers* and *Probability-of-Dissociation-Trimers* (98%, i.e., our standard condition, and 50%) highlight the pivotal role of trimers in determining the timing for the start of the elongation phase. A marked decrease in dimers and trimers instability may be interpreted as an overexpression of nucleators or an enhancement of their activity. Based on our observations, we fixed the simultaneous presence of 5 polymers as the arbitrary threshold above which elongation is irreversible (i.e., the elongation phase has started). When both probabilities of dissociation of dimers and trimers are 98%, elongation is reached 53 ticks after the start of the simulation. This time interval slightly decreases when the probability of dissociation of dimers is reduced at 50% (49 ticks). Instead, the effect of a reduction in the probability of dissociation of trimers (50%) is way more pronounced, leading to the start of elongation in 22 ticks, which confirms the importance of nucleating dynamics and the key role of trimers in the actin polymerization system.

### Evolution of filament lengths distribution and steady state

Numeric data for filament lengths distribution were obtained from the histograms generated during a single NetLogo run at different time points referring to both the extension and steady state phases (i.e., 100, 200, 300, 400, 600, 800, 1,000, 1,500, 2,000 and 5,000 ticks). Using these data, we first constructed the corresponding density plots (shown in Fig. 5). These curves show that, as the simulation proceeds, the number of length classes gradually increases until a slow phase where monomers are redistributed among the filaments is reached. This phase is characterized by the dominance of the so-called “diffusive regimen” [12].

**Fig. 5.**
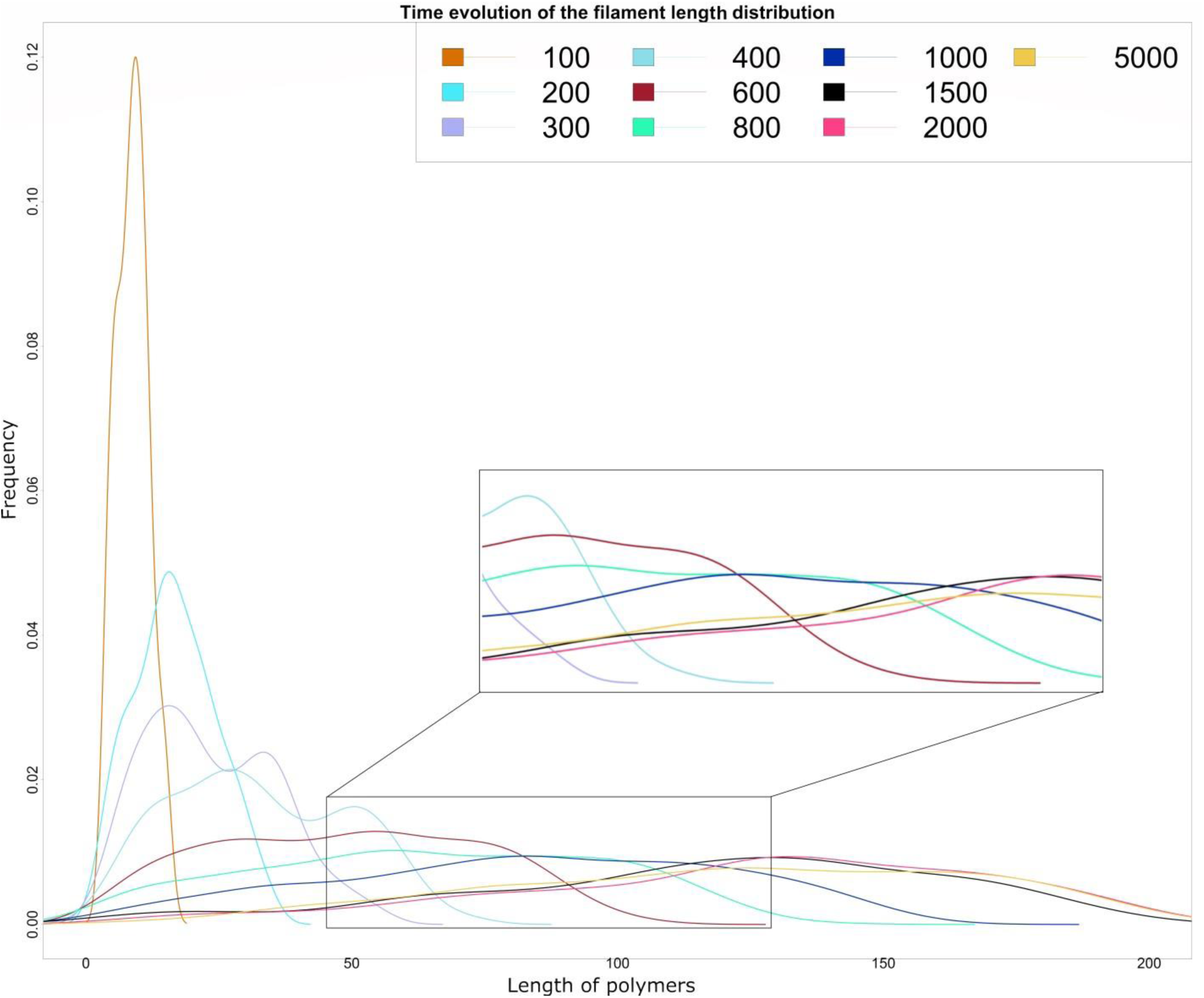
Evolution of length distribution over time. Times in ticks and colors of the corresponding curves are shown in the box at the top right corner.

To examine how polymer lengths evolve over time, we analyzed NetLogo length distributions of filaments at different time points of a single run (i.e., 6000, 9000, 15000, 24000, 31000, 4300, 52000, 57000, 70000, 81000, 87000, 106000, and 125000) by running pair-wise comparisons between those distributions (see Table 4 and Supplemental File 1). Significant differences only appear when the time intervals between the considered ticks exceed 19,000 ticks, suggesting that polymer lengths are weakly stable over time.

**Table 4.** Kolmogorov-Smirnov p-values from pair-wise distribution comparisons.

| Ticks.time.1 | Ticks.time.2 | K-S.pvalues | K-S.sign<sup>^</sup> |
| --- | --- | --- | --- |
| 6000 | 9000 | 0,9 |  |
| 6000 | 15000 | 0,453 |  |
| 6000 | 24000 | 0,154 |  |
| 6000 | 31000 | 0,0929 |  |
| 6000 | 43000 | 0,00456 | ** |
| 6000 | 52000 | 0,000726 | *** |
| 6000 | 57000 | 0,00117 | ** |
| 6000 | 70000 | 0,00445 | ** |
| 6000 | 81000 | 0,00114 | ** |
| 6000 | 87000 | 0,00101 | ** |
| 6000 | 106000 | 1,43E-08 | **** |
| 6000 | 125000 | 7,52E-09 | **** |
| 9000 | 15000 | 0,681 |  |
| 9000 | 24000 | 0,272 |  |
| 9000 | 31000 | 0,135 |  |
| 9000 | 43000 | 0,00802 | ** |
| 9000 | 52000 | 0,00211 | ** |
| 9000 | 57000 | 0,00381 | ** |
| 9000 | 70000 | 0,00801 | ** |
| 9000 | 81000 | 0,00259 | ** |
| 9000 | 87000 | 0,00162 | ** |
| 9000 | 106000 | 1,02E-07 | **** |
| 9000 | 125000 | 2,56E-08 | **** |
| 15000 | 24000 | 0,826 |  |
| 15000 | 31000 | 0,533 |  |
| 15000 | 43000 | 0,0899 |  |
| 15000 | 52000 | 0,0168 | * |
| 15000 | 57000 | 0,0317 | * |
| 15000 | 70000 | 0,0471 | * |
| 15000 | 81000 | 0,0349 | * |
| 15000 | 87000 | 0,0162 | * |
| 15000 | 106000 | 2,89E-06 | **** |
| 15000 | 125000 | 8,19E-07 | **** |
| 24000 | 31000 | 0,864 |  |
| 24000 | 43000 | 0,507 |  |
| 24000 | 52000 | 0,118 |  |
| 24000 | 57000 | 0,273 |  |
| 24000 | 70000 | 0,17 |  |
| 24000 | 81000 | 0,114 |  |
| 24000 | 87000 | 0,0919 |  |
| 24000 | 106000 | 0,000234 | *** |
| 24000 | 125000 | 1,83E-05 | **** |
| 31000 | 43000 | 0,907 |  |
| 31000 | 52000 | 0,379 |  |
| 31000 | 57000 | 0,299 |  |
| 31000 | 70000 | 0,408 |  |
| 31000 | 81000 | 0,242 |  |
| 31000 | 87000 | 0,203 |  |
| 31000 | 106000 | 0,000205 | *** |
| 31000 | 125000 | 4,53E-05 | **** |
| 43000 | 52000 | 0,557 |  |
| 43000 | 57000 | 0,688 |  |
| 43000 | 70000 | 0,592 |  |
| 43000 | 81000 | 0,732 |  |
| 43000 | 87000 | 0,452 |  |
| 43000 | 106000 | 0,00161 | ** |
| 43000 | 125000 | 0,000235 | *** |
| 52000 | 57000 | 0,986 |  |
| 52000 | 70000 | 0,943 |  |
| 52000 | 81000 | 0,812 |  |
| 52000 | 87000 | 0,688 |  |
| 52000 | 106000 | 0,0161 | * |
| 52000 | 125000 | 0,00421 | ** |
| 57000 | 70000 | 0,994 |  |
| 57000 | 81000 | 0,804 |  |
| 57000 | 87000 | 0,936 |  |
| 57000 | 106000 | 0,0575 |  |
| 57000 | 125000 | 0,013 | * |
| 70000 | 81000 | 0,894 |  |
| 70000 | 87000 | 0,926 |  |
| 70000 | 106000 | 0,0128 | * |
| 70000 | 125000 | 0,00498 | ** |
| 81000 | 87000 | 0,971 |  |
| 81000 | 106000 | 0,0346 | * |
| 81000 | 125000 | 0,00524 | ** |
| 87000 | 106000 | 0,035 | * |
| 87000 | 125000 | 0,0103 | * |
| 106000 | 125000 | 0,988 |  |

^ sign. p-values: \* = $p < 0.05$ ; \*\* = $p < 0.01$ ; \*\*\* = $p < 0.001$ ; \*\*\*\* = $p < 0.0001$ .

### Treadmilling

In the steady state phase, the polymerization dynamics enter an equilibrium state (diffusive regimen) in which the dissociation of monomers from the pointed-end (-) and the association to the barbed-end (+) are balanced and maintained by a critical concentration of free monomers in the cytosol. This steady state of assembly and disassembly is known as treadmilling. We found that a key feature of our NetLogo model is its ability to successfully reproduce this important molecular phenomenon. Indeed, to detect the presence of treadmilling during simulations, we used the following criteria:

1. the ratio between monomers association rate and monomers dissociation rate must be = 1 (see Fig. 6);
2. the simulation must be in the steady state, a condition in which the number of polymers does not change, or is nearly constant over time;
3. at the plateau, the number of free monomers must remain constant or nearly constant within a given range of close values.

**Fig. 6.**
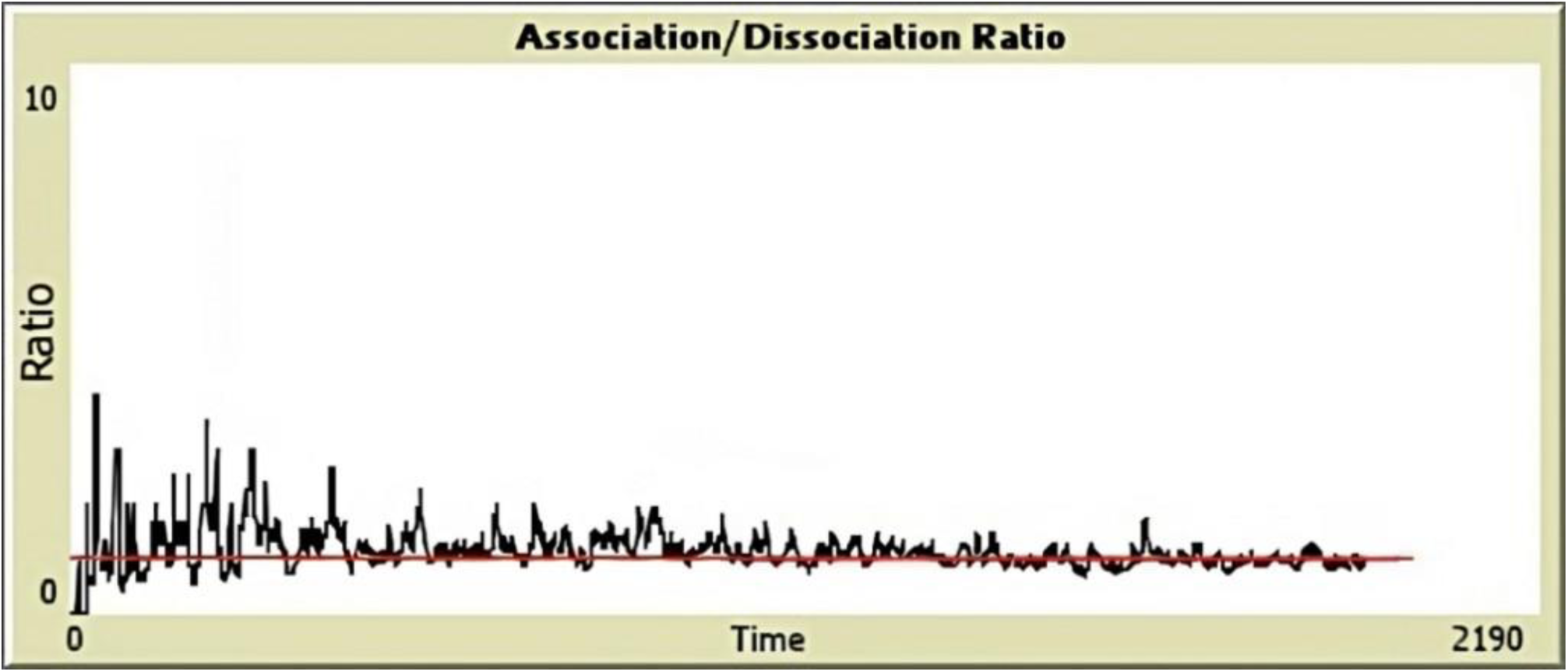
Monomer association/dissociation ratio over time. Plot of the model’s interface displaying the ratio of monomer association (barbed-end) and dissociation (pointed-end) rates. A value approximating to a ratio = 1 (represented by the red line) suggests the occurrence of treadmilling.

Hereafter, we illustrate a specific case study documenting how treadmilling can be detected during a simulation with the NetLogo model. If the following initial conditions are used: 5,000 *Initial-Monomers*, *Probability-of-Dissociation-Polymers* = 8%, *Probability-of-Dissociation-Dimers* = 98%, and *Probability-of-Dissociation-Trimers* = 98%, treadmilling is reached at about 1,200 ticks and can be identified: (i) when the value = 1 appears in the monitor displaying the ratio between association and dissociation rates and/or (ii) when in the *V_ass_/V_diss_* plot the ratio value curve (in black) overlaps the red line that corresponds precisely to the value of 1 (see Fig. 6).

We also investigated the role of dimers in the maintenance of both the steady state and the ratio between association and dissociation rate = 1. In these runs, we introduced an abrupt change in the probability of dissociation of dimers during the steady state. More specifically, we reduced the value of the parameter *Probability-of-Dissociation-Dimers* from our standard 98% to 30% while the system was in the steady state (*Initial-Monomers* = 15,000, *Probability-of-Dissociation-Trimers* = 98%, *Probability-of-Dissociation-Polymers* = 8%). This is a speculative scenario, since it is well-acknowledged that dimers are not more stable than trimers [36]. As expected, we found that decreasing the degree of dimers instability results in the disruption of the previously established steady state (indeed, this change induces a new phase of growth in the number of polymers, see Fig. 7). Simultaneously, the association/dissociation ratio returns almost constantly, although slightly greater than 1 (Fig. 8). These outcomes seem to strengthen our point that having an association/dissociation ratio = 1 may be considered a diagnostic criterion to detect actin treadmilling.

**Fig. 7.**
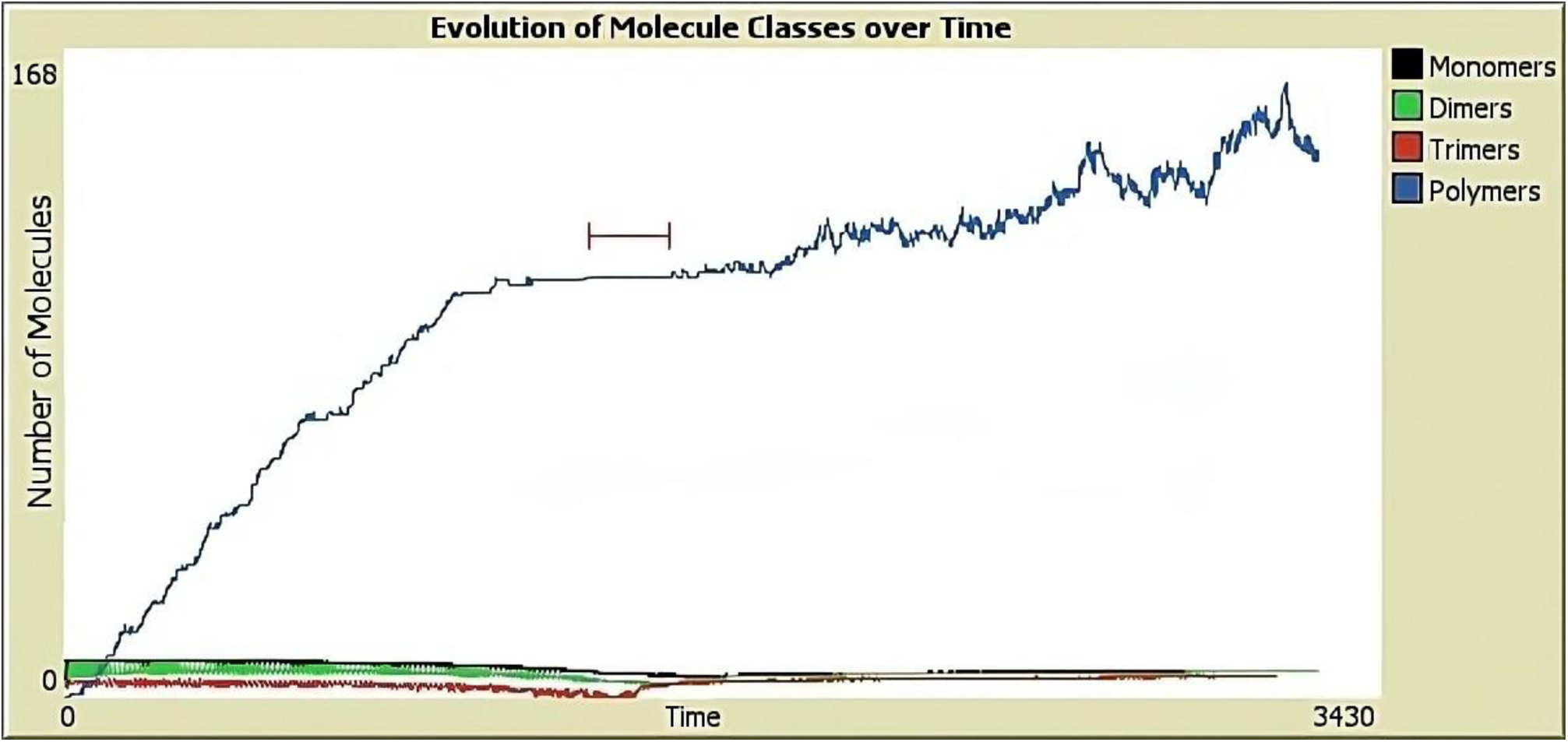
Effects of decreasing *Probability-of-Dissociation-Dimers* on the evolution of molecule classes over time. After an interval of ticks (1360-1585) characterized by treadmilling (i.e., the steady state phase, see black interval in the figure), the increased stability of dimers led to the start of a new elongation phase. Note that, in this figure, we used a logarithmic scale to represent the curves of monomers, dimers, and trimers, and a non-logarithmic scale to represent polymers.

**Fig. 8.**
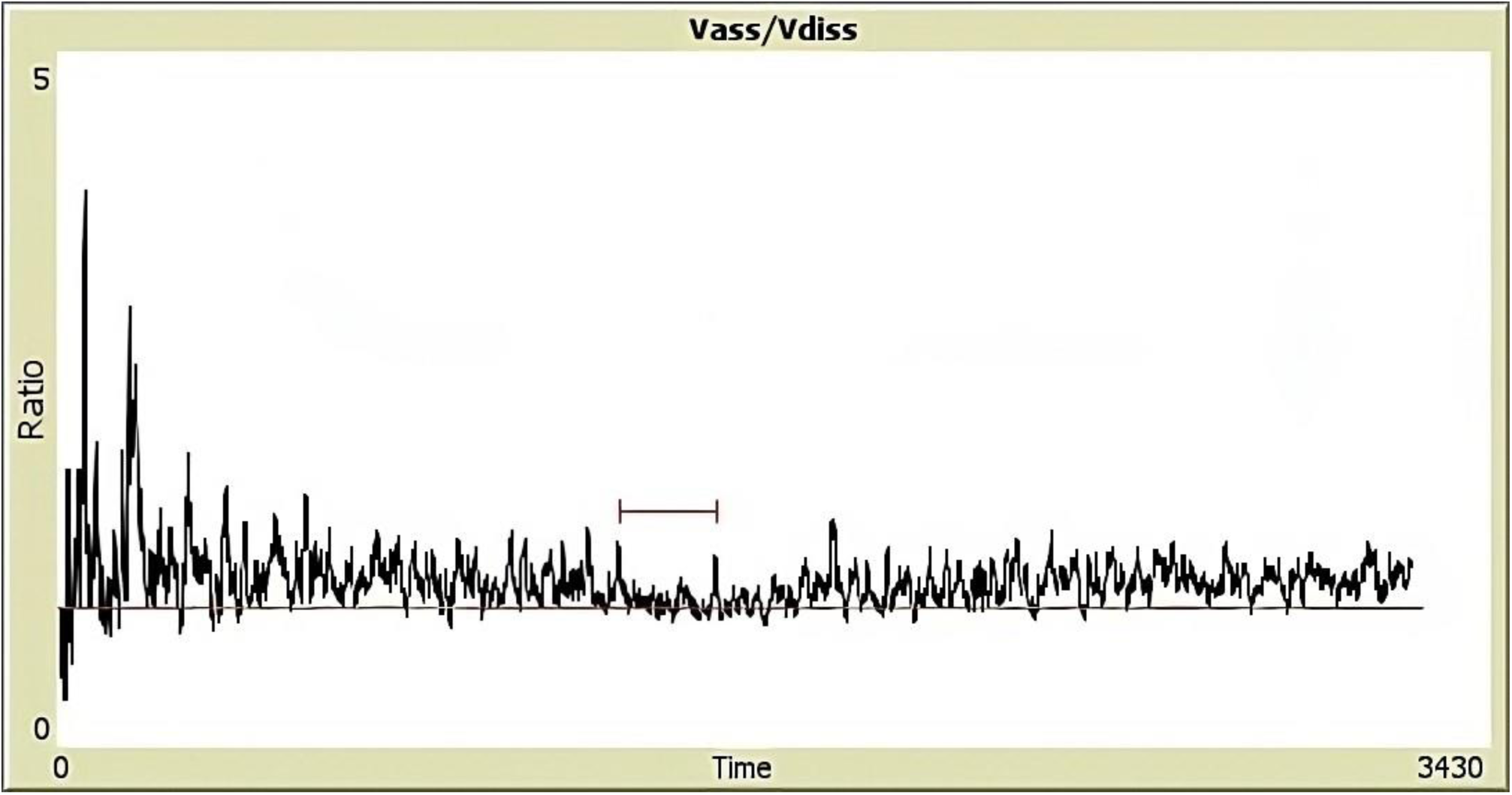
Effects of decreasing *Probability-of-Dissociation-Dimers* on the association/dissociation ratio. After an interval of ticks (1360-1585) characterized by treadmilling (black interval, where V_ass_/V_diss_ = 1.04, SD = 0.08), the increased stability of dimers led to the start of a new elongation phase (V_ass_/V_diss_ = 1.17, SD = 1.12).

### Competition between growth in the number of polymers and the average length of polymers

Simulations were also carried out to observe how the average length of polymers varies as a function of the number of initial monomers. For a given number of initial monomers, two competing processes are expected to take place: (i) the elongation of polymers which are already present in the simulation environment by net association of new monomers and (ii) the formation of new polymers from the aggregation of monomers and trimers to form tetramers (i.e., the shortest polymers). Both these processes are fed by the same pool of initial monomers. A parametric analysis (summarized in Table 5) was thus carried out by varying the value of *Initial-Monomers*, in order to investigate the effects of this parameter on the two competing processes. To analytically assess the hypothesis of competition between the growth rates of P (number of polymers) and L_m_ (average polymers length), we analyzed the relationship between these two variables using transient plots in NetLogo, where P is the ordinate and L_m_ is the abscissa, for several values of initial monomers (i.e., from 1,000 up to 40,000). For each number of initial monomers, we first exported numerical values of P and L_m_ at t = 500 ticks and tested their correlation with the cor.test () function in R. The selected time corresponds – for all considered cases – to the extension phase of polymerization. In all cases we obtained significant correlation values, with r > 0.95. Then we calculated the averages of P and L_m_, here denoted as 𝑃̅ and L̅̅_m̅̅_ respectively, and their ratio. This ratio could be interpreted as the slope of the hypothetical regression line constructed on each scatterplot of P vs L_m_.

**Table 5.** Competition between the synthesis of new polymers and elongation of existing polymers in terms of P/L_m_ slopes.

| *Initial-Monomers* | *Probability-of-Dissociation-Polymers = 8%* |  |  |
| --- | --- | --- | --- |
| | $\overline{L}_m$ | $\overline{P}$ | $\overline{P}/\overline{L}_m$ |
| 1000 | 30 | 3 | 0.10 |
| 2000 | 28 | 4.8 | 0.17 |
| 3000 | 25 | 6 | 0.24 |
| 4000 | 25.5 | 11.8 | 0.46 |
| 6000 | 24.1 | 14.1 | 0.58 |
| 8000 | 22.2 | 18.3 | 0.82 |
| 10000 | 21.8 | 22.9 | 1.05 |
| 15000 | 22.8 | 27.5 | 1.20 |
| 20000 | 23.6 | 44.9 | 1.90 |
| 25000 | 23.2 | 49.7 | 2.14 |
| 30000 | 23 | 60 | 2.60 |
| 35000 | 23.9 | 82.3 | 3.44 |
| 40000 | 22.1 | 78.7 | 3.56 |

\*Data were extracted at 500 ticks, i.e., during the extension phase.

In this context, Fig. 9 displays the aforementioned procedures for two representative values of the initial number of monomers, i.e., 2,000 and 35,000.

**Fig. 9.**
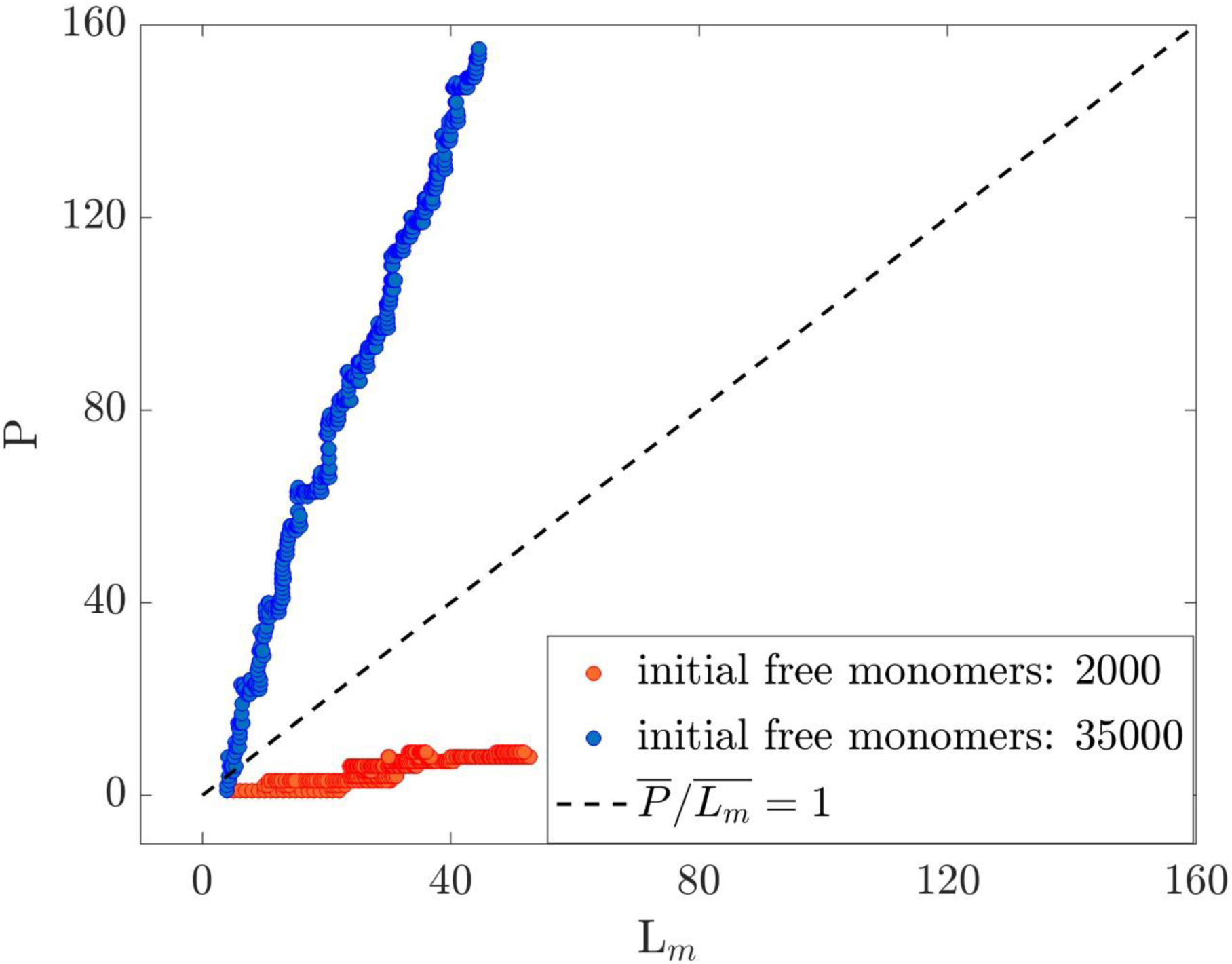
Time evolution of polymer lengths in monomeric subunits (abscissa) vs number of polymers (ordinate) for two selected values of the initial number of free monomers. In this plot, red symbols correspond to an initial number of 2,000 free monomers, while blue symbols correspond to 35,000 free monomers. Also, the discontinuous black line in the graph reports the unitary slope, denoting how a higher number of free initial monomers promotes the formation of new polymers with respect to the elongation of the existing ones observed at lower concentrations.

Furthermore, the slope values obtained for a wide range of initial monomers (i.e., from 1,000 to 40,000) were plotted against the initial number of monomers to obtain the graph in Fig. 10. This figure informs that, for low values of initial monomers (from 1,000 to 10,000), the growth rates of polymer lengths are higher than the growth rates of the number of polymers (see values below the red line, whose ordinate is fixed = 1). In contrast, the rates above the red line (slopes > 1) correspond to simulations carried out with initial monomer values > 10,000. In the latter case, the growth rate of the number of polymers outweighs that of lengths. The analysis clearly shows a dependence of P and L_m_ growth rates on the initial number of monomers, which can be rephrased as follows: few initial monomers tend to form long polymers, while many initial monomers tend to form many, but short polymers.

**Fig. 10.**
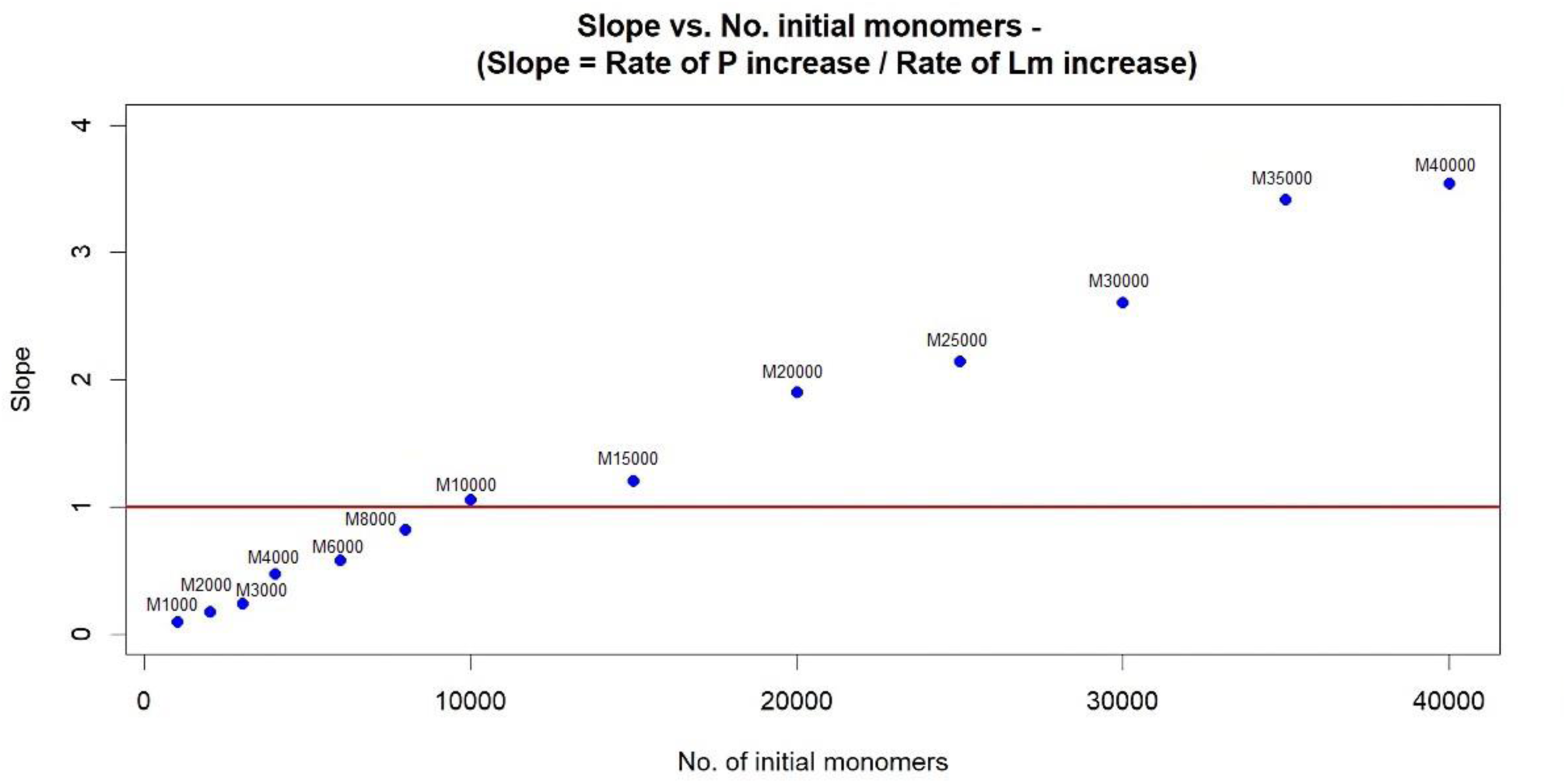
Competition between the increase in number and in the average length of polymers. Growth rates of average polymer lengths are greatest for an initial number of monomers < 10,000 (see values of slopes below the red line). Conversely, for values of initial monomers > 10,000, the growth rates of the number of polymers prevail (see also Table 3). L̅̅_m̅̅_ = average length of filaments; 𝑃̅ = average number of polymers.

The ability of the model to reproduce the previously described competition was further confirmed by displaying the changes in L_m_ and P at convergence (i.e., at the beginning of the steady state) as a function of the number of initial monomers, as shown in Fig. 11. In a few words, the numerical data corresponding to the start of the steady state were exported from L_m_ vs P plot for each simulation performed with a particular number of initial monomers. These values of L_m_ and P at convergence were then plotted as a function of the number of initial monomers. As can be seen from Fig. 11, for values below 10,000 initial monomers the increase in length prevails (see abrupt growth of the blue curve), whereas for values above 10,000 initial monomers the lengths remain constant (see blue curve). In contrast, the curve of the number of polymers plotted as a function of the initial number of monomers shows a linear increase in the number of polymers that continues even for values of initial monomers greater than 10,000 (see red curve).

**Fig. 11.**
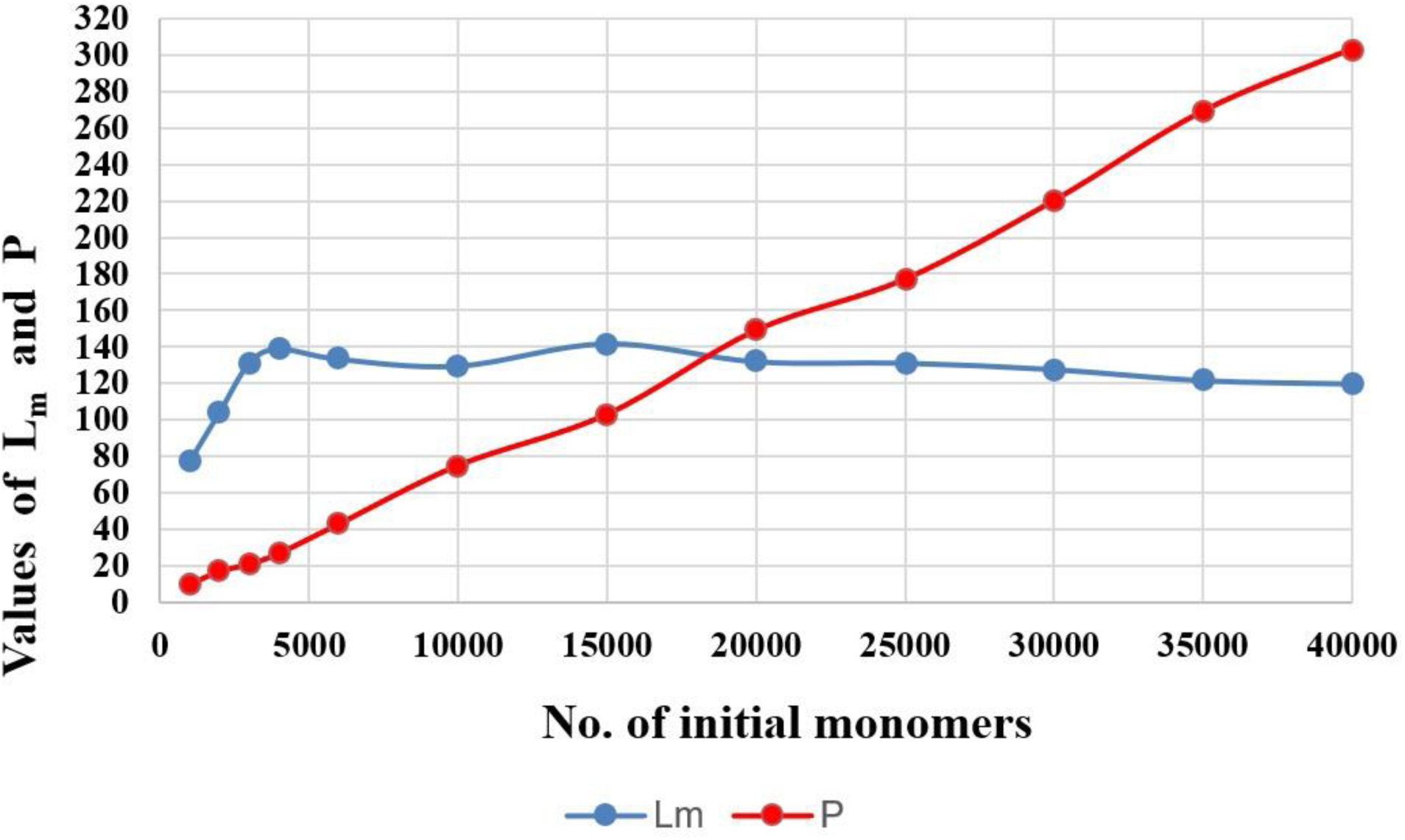
Different growth rates for mean length (L_m_) and number of polymers (P) depend on the initial number of monomers.

Collectively, the results of these three analyses support the existence of a competition between the increase in the number of polymers and the increase in polymer lengths. Up to 10,000 initial monomers, the increase in L_m_ prevails, while for higher values of initial monomers the prevailing process is the synthesis of new polymers.

### The intensity of actin polymerization is correlated to the increase in the length of polymers

Another outcome of relevance was obtained when focusing on the relationship between the overall behavior of the simulated system and our indicator of the degree of polymerization, that is, the Carothers’ equation. More specifically, between the end of the extension phase and the start of the steady state (at convergence), we observed an abrupt increase in the degree of actin polymerization as calculated by the equation of Carothers. This effect appears to be more correlated with an increase of the average length of polymers (r = 0.76), rather than an increase of the number of polymers (r = 0.56). A possible interpretation of this result is given in the Discussion.

## Discussion

In this study, we developed a NetLogo model of the actin polymerization process in vitro, that is, mimicking a cell-free system based on G-actin monomers and including the effects of some other proteins that, in vivo, are usually implicated in the process (e.g., Cofilin). Our main objective was to provide a digital environment suitable for replicating the temporal evolution of the system, emphasizing the coexistence of populations of monomers, oligomers, and polymers.

Several studies cited in the Introduction investigated various aspects of actin involvement in various cellular or subcellular contexts. Although some of these studies also used agent-based modeling (e.g., [18]), to our knowledge, this is the first ABM implemented with NetLogo that successfully reproduces the whole process of actin polymerization, including some emerging, higher-level patterns which are spontaneously generated by the system, such as treadmilling and competition between nucleation and elongation. Consequently, the model is validated by the strong qualitative correspondence between the outcomes of simulations and the real phenomena observed in vitro. This makes the “Actin Polymerization” NetLogo model a reliable platform to support empirical research, for instance, by allowing scientists to explore molecular dynamics or speculative scenarios at the basic level or even by suggesting new hypotheses.

In this regard, we want to recall the main similarities of the model we developed with respect to what is known about the in vitro process. As Table 2 shows, the design of our model included seven out of fifteen features considered for the phenomenon of actin polymerization. The simulations presented in this article confirm that the limited set of features implemented into the model are sufficient to reproduce the three main phases of polymerization, that is, nucleation, extension, and steady state. As shown by our analyses, the overall system’s dynamics reflect the real behavior of each step of this complex molecular mechanism. In particular, by simulating the entire polymerization process, it was possible to investigate the time evolution of the distribution of filament lengths. This analysis showed that, as the simulation proceeds, a progressive increase in the number of polymer length classes is observed. In agreement with Fujiwara et al. [13] and Hu et al., [10], our investigations into length distribution conducted at the steady state showed that polymerization/depolymerization dynamics follow a diffusion (stochastic) process, which cannot be explained by the simple association and dissociation of monomers at the two ends of the filaments respectively. Interestingly, our Kolmogorov-Smirnov statistics revealed that the steady state phases are transient or intermittent. In particular, these phases can be observed even for long time intervals when comparing length distributions from two closer time points (e.g., 24,000 and 87,000 ticks, see Table 4), whereas they are disrupted when comparing distributions from two more distant time points (e.g., 31,000 and 125,000 ticks). Although it would be necessary to conduct further investigations to determine whether the long-term behavior of the system could represent a real biological phenomenon (which is plausible, since even a steady state limited in time might be persistent enough to enable cells to carry out their functions effectively), we can conclude that our model is able to reproduce the actin steady state. Its time limitations may also be due to the absence of regulatory proteins and mechanisms that we did not include in our model but that could have long-term effects on the behavior of the system, or to the absence of a regulable probability of association of monomers to the barbed ends (see also the subsection below dedicated to the limitations of the model).

Above all, in the steady state phase, it was possible to observe a spontaneous global behavior which largely corresponds to the molecular phenomenon of treadmilling (see diagnostic criteria used in the “Treadmilling” subsection of the Results). This outcome is particularly relevant since no procedure for generating treadmilling at a system level was implemented in the model’s code, in line with the core principles of agent-based modeling. Thus, treadmilling appears as an emergent property of the actin polymerization complex system. Furthermore, we showed that treadmilling occurs on coexisting subpopulations of filaments, with a constantly high degree of synchronization of association and dissociation over time for most polymers. This phenomenon is analogous to the global treadmilling described by several authors [37,38]. The most notable difference between the two systems is that, in our case, treadmilling emerges as an *intrinsic* property of actin polymerization, because our computational model does not include all features of the in vitro environment which were considered, for instance, in the study of Carlier and Shekhar [37].

Another result also dealing with the emerging patterns of actin polymerization was the identification and analysis of the competition between the formation of new polymers and the elongation of pre-existing ones, a competition that – as we have shown – is highly dependent on the initial number of monomers which are present in the system. Indeed, for example, if a simulation is run with a few hundred initial monomers, only a few polymers are obtained, but they are long. In contrast, a simulation conducted with many initial monomers (> 1,000) tends to generate many, but shorter polymers. We further investigated the competition between new polymer synthesis and the elongation of existing polymers considering the degree of polymerization resulting from the Carothers’ equation. We observed that, near the end of the extension phase and at the beginning of the steady state, there is a significant acceleration of the degree of polymerization. In principle, the degree or intensity of polymerization may depend on an increase in the number of new polymers, as well as an increase in the length of existing polymers. In this regard, we found a higher correlation with an increase in the length of polymers. This higher correlation can be interpreted as an indicator of competition between the synthesis of new polymers and the process of elongation. In summary, we obtained two evidences of competition occurring at different stages of polymerization: (i) at the start of polymerization, when the first oligomers are formed, and (ii), at the final stage of extension/beginning of the steady state. This competing behavior is consistent with recent in vitro and theoretical findings [21,22,39,40]. In particular, Zweifel et al. [21] studied the role that Formin plays in the process of nucleation and filament extension in vitro. They found that these two processes influence each other, as the monomeric pool from which they draw is the same. This interdependence determines the number of filaments assembled during a polymerization reaction, as well as their lengths. Furthermore, these authors observed that the speed of the Formin-mediated polymerization reaction appears to depend on the concentration of available actin monomers: as the concentration of actin monomers increases, the number of assembled filaments increases, but the average length decreases. Finally, in their work, Zweifel et al. [21] emphasized the advantage of keeping the nucleation efficiency of Formin low in cells to prevent dysregulated actin assembly, which would have deleterious consequences on the dynamics of the cell’s cytoskeleton. In our model, the activity of Formin could simply correspond to the instruction in the code determining the binding of two agents (i.e., monomer + barbed-end) following co-localization in the same patch. Consequently, it seems to us of relevance that the competition can be detected in such a simplified computational system. Together with treadmilling, this high-level behavior emerges as a global property of the system.

Collectively, the simulations performed with NetLogo showed that the dynamics of actin polymerization are, in many ways, comparable to the typical features of complex systems. Table 6 shows the main correspondences between an “ideal” complex system and the actin polymerization process simulated in NetLogo. Moreover, Fig. 12 shows all interactions and feedback loops involved in the model.

**Table 6.**
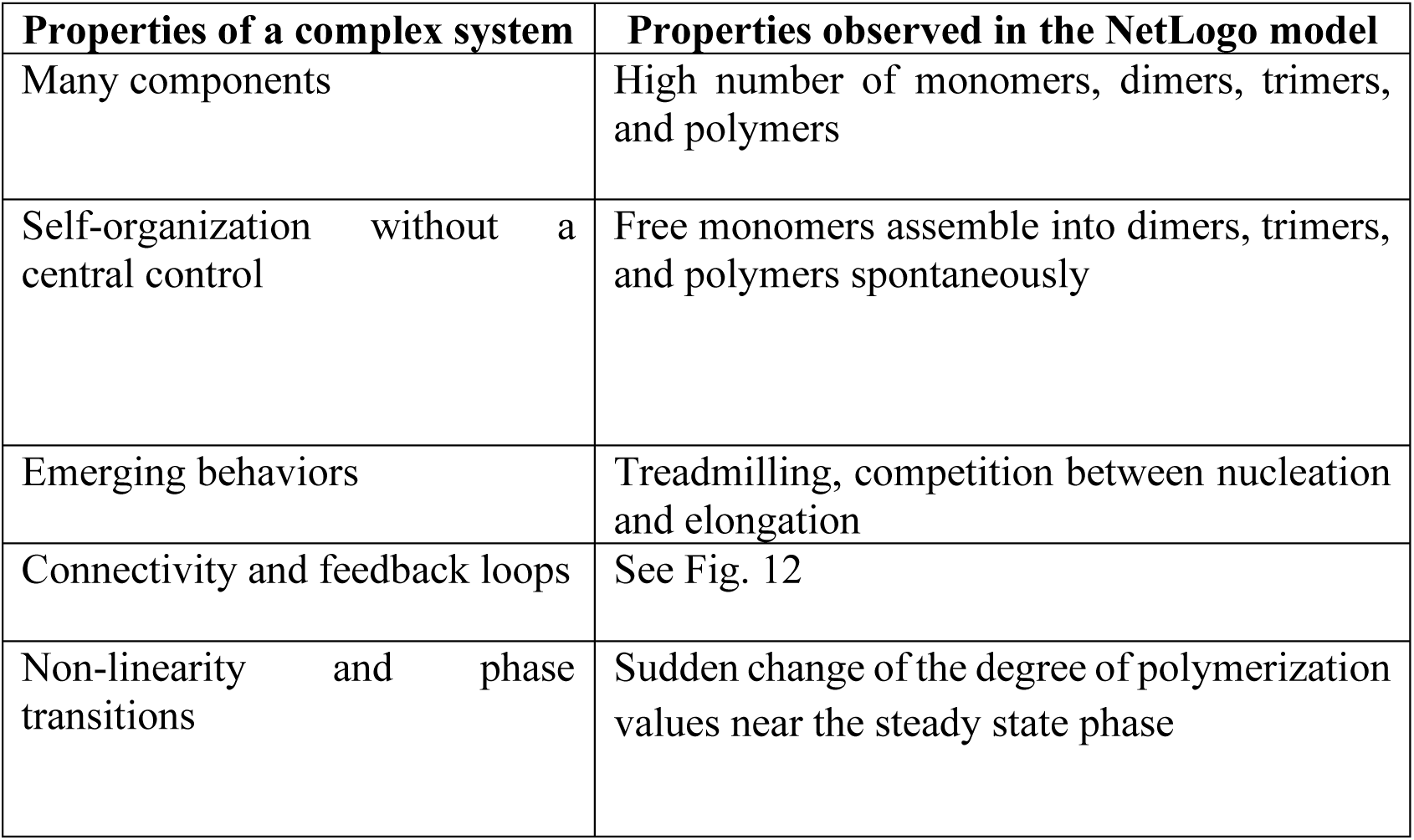
Main properties a typical complex system shares with the actin polymerization model developed in NetLogo.

**Fig. 12.**
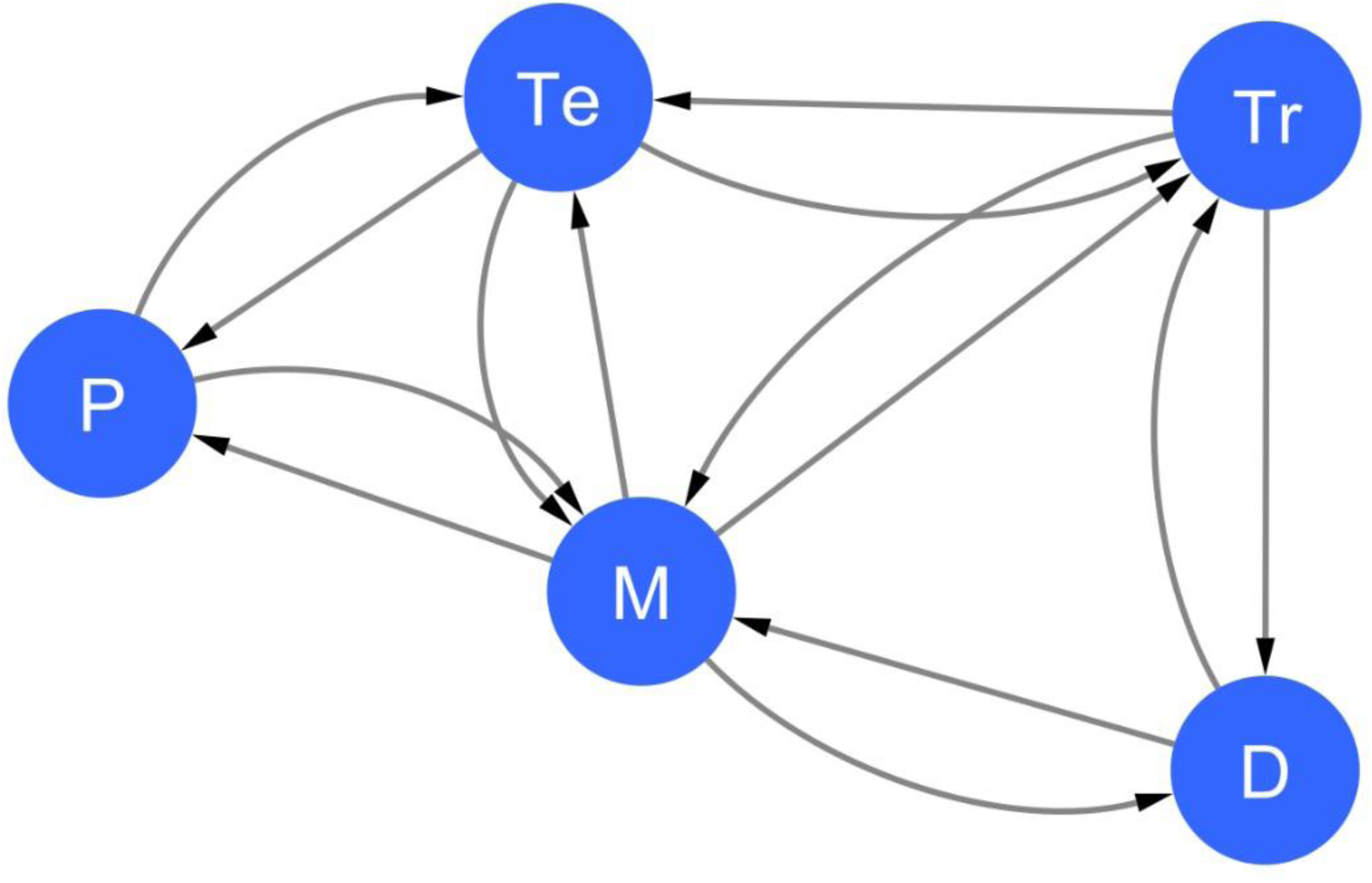
Network representing all interactions involved in the model. Network including the interactions between monomers (M), dimers (D), trimers (Tr), tetramers (Te) and polymers (P) during the actin polymerization process. The M node has the highest degree (hub) because it has the highest number of links (8). Note the large number of feedback loops.

### Limitations of the model

We have already mentioned several features distinguishing our model from the experimental (in vitro/in vivo) actin system, e.g., in Table 2. Indeed, though our objective was to model actin polymerization and its emerging dynamics, we were only partially able to achieve this goal. Like any in silico modeling of biomolecular processes, our new NetLogo model also has some limitations that could circumscribe the validity of some outcomes to very specific contexts. On the other hand, many of the patterns described so far could be applicable as well to other polymer chains that can grow and shrink from both ends (e.g., tubulin). Here, we focus on some of the main limitations of the current version of the model.

The first one concerns the lack of experimental connection to the parameter value chosen for the probability of dissociation of trimers. Hereafter, we provide the rationale behind the choice of probability of 98% for both dimers and trimers. Calculation based on dissociation rates listed in [8] suggests that, if we consider a 98% probability of dissociation of dimers, then the corresponding probability of dissociation of trimers should be around 0.04%. However, using these probabilities (i.e., approximating 0.04% to 1% for trimer dissociation), simulations still generate all three phases of polymerization (i.e., nucleation, elongation, and steady state). The main variation observed when using these values is a reduction in the mean length of polymers. If on one hand the high dissociation probability for trimers appears very different from experimental evidence, on the other hand, there are hints from the literature supporting our choice. In the first place, in the context of spontaneous polymerization of actin monomers, the first two steps of nucleation (i.e., formation of dimers and formation of trimers) are both very unfavorable, with very high instability of both oligomers, whereas greater stability is only achieved with tetramers [8]. So, one of the reasons why we used a very high probability of dissociation for both dimers and trimers was the necessity of simulating the effects of extremely unstable oligomers below 4 subunits (i.e., below tetramers). In this regard, we recall that all our runs were executed considering a system where nucleators are absent or show a very low activity, so one can focus only on the emerging behavior generated by a limiting monomer pool. These conditions, which are close to those considered in some recent studies, support the assumption of a high dissociation rate for both dimers *and* trimers [22]. In the real world, our parameter values could reflect in vitro/in vivo conditions characterized by reduced functioning of the nucleation system, which would result in a very low number of stable trimers (e.g., see [41]).

The second issue involves the absence from the code of procedures which implement several mechanisms which could influence the global dynamics of the system, including capping, severing, and annealing. In particular, annealing of filaments, that is, the process where two shorter actin filaments join end-to-end to form a longer filament, could affect the competition between filament nucleation and growth. However, consistently with our assumptions, we share the same observation proposed in [22] where authors point out that while “filament turnover by spontaneous fragmentation, annealing, and cofilin-mediated severing may affect the long-time actin dynamics in an actin concentration-dependent manner, thereby altering the time evolution of filament length distribution […] we do not consider such effects and restrict our study to smaller actin concentrations, focusing on short timescales compared with the timescales over which fragmentation or annealing effects become prominent”.

The third issue concerns the correspondence between the time of simulations in ticks and the time in seconds required for biochemical processes. Although in principle such correspondence could be established in relative terms (e.g., using proportions), this approach was not included in our study, since our emphasis focused only on simulation time. For this reason, this and other aspects concerning the comparability between our simulated time intervals and real biochemical time intervals would need further investigation in future assessments.

Finally, as shown in the Results, steady state appears to be a transient phenomenon followed in the long-term by phases where filament lengths increase. Although this could reflect real biological dynamics, such an effect may also be due to the absence of procedures implementing the activity of several regulatory proteins controlling the various phases of polymerization.

## Conclusions

Overall, the performance of the model appears satisfactory, suggesting that the selection of features from the in vitro process implemented in the algorithm was done properly. However, the above limitations and the absence of an accurate parameterization make our simulated scenarios less specific for the actin system and lead us to think of the model as a toy model that could be applied to investigate or illustrate the dynamics of other biomolecular systems sharing the main functional assumptions considered in this work. The model can be improved by including new characteristics that can further reduce the number of dissimilarities with an in vitro system. The features that we plan to add include a function controlling the probability of attaching monomers to the barbed-end and the possibility of regulating the association and dissociation rates of monomers at both ends of filaments. A further improvement could also be the development of a 3D version of the model. In its present version, the model can be used to test the effects of combining several parameters on actin polymerization dynamics. Therefore, this model may contribute to shed light on yet poorly understood aspects of the whole actin polymerization process.

## Acknowledgements

The authors wish to thank Prof. Diego Molteni and Prof. Renato Lombardo of the University of Palermo for their valuable suggestions and helpful discussion.

## Data availability statement

The NetLogo model we implemented and used for our experiments can be freely downloaded from the following link: https://doi.org/10.5281/zenodo.17860349. The model can be opened by installing the NetLogo simulation platform (version 6.2.0), which can be downloaded free of charge from the official website of the software at the following link: https://ccl.northwestern.edu/netlogo/6.2.0/.

## Author contributions

R.T.: Conceptualization, Data Curation, Investigation, Methodology, Software, Validation, Visualization, Writing – original draft, Writing – review and editing

S.C.: Conceptualization, Data Curation, Formal Analysis, Investigation, Validation, Visualization, Writing – original draft, Writing – review and editing

L.G.: Data Curation, Investigation, Validation, Visualization, Writing – original draft, Writing – review and editing

G.I.: Conceptualization, Data Curation, Investigation, Formal Analysis, Validation, Visualization, Writing – original draft, Writing – review and editing

G.P.: Data Curation, Formal Analysis, Visualization, Writing – original draft, Writing – review and editing

G.B.: Data Curation, Formal Analysis, Visualization, Writing – original draft, Writing – review and editing

V.R.: Conceptualization, Investigation, Supervision, Validation, Writing – original draft, Writing – review and editing

## Supplemental material

**Supplemental file 1. ECDFs showing the pair-wise comparisons between length distributions.**

