## Supplemental Figure 1 for "An Agent-Based Model of Protein Polymerization Dynamics: Focus on the Actin System"

**Supplemental File 1**

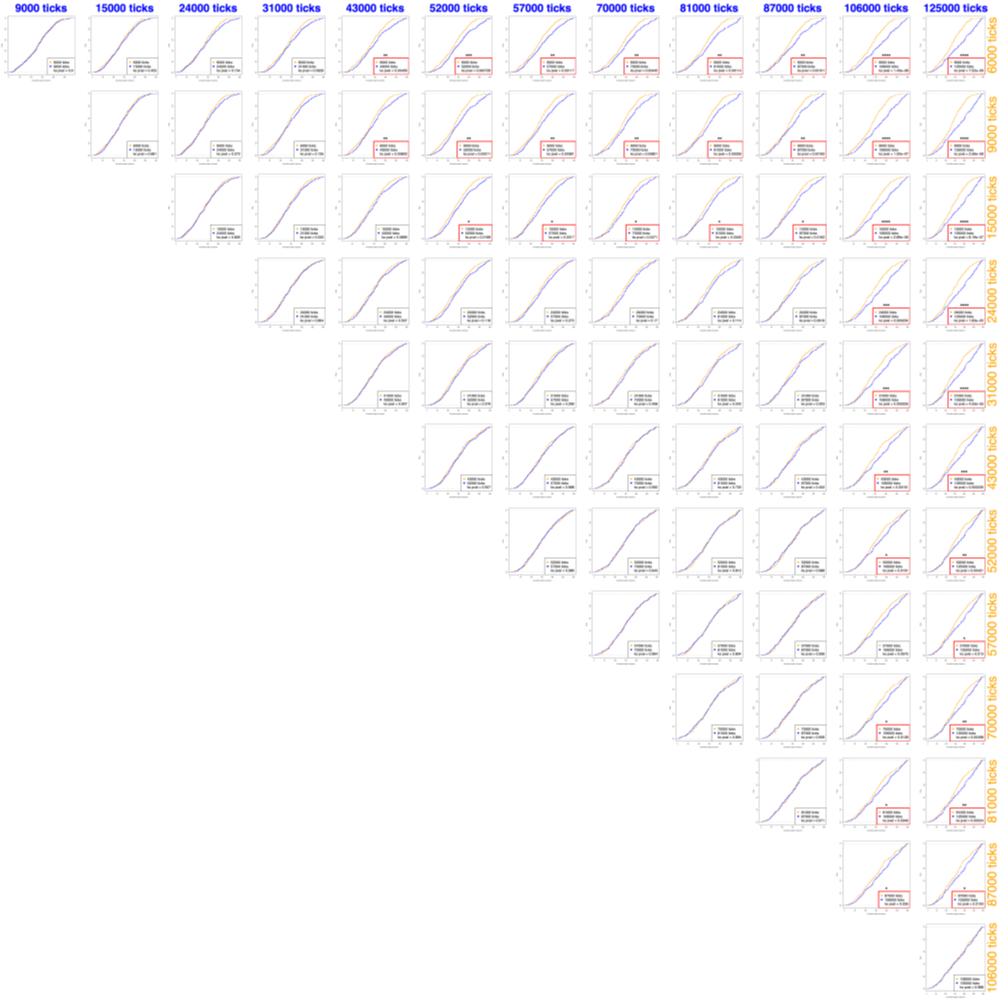

**Supplemental Figure 1. ECDFs showing the pair-wise comparisons between length distributions.**
